# Effect Of Perinatal Perfluorinated Compounds (PFASs) Exposure On Behavior Of SD Rats

**DOI:** 10.1101/2025.03.16.643539

**Authors:** Yuzi Sun, Dingyue Chai, Jiamin Lu, Yuhui Yao, Qianqian Cai, Min Peng, Lihui Wu

## Abstract

Perfluorinated compounds (PFASs) are persistent environmental pollutants recognized for their bioaccumulation and potential neurotoxic effects, particularly in relation to attention deficit hyperactivity disorder (ADHD). This study investigated the effects of perinatal PFAS exposure on Sprague-Dawley (SD) rats, with a focus on behavioral outcomes and dopamine metabolism in the prefrontal cortex (PFC). Pregnant rats were administered PFASs at concentrations of 1 mg/L, 3 mg/L, and 10 mg/L through drinking water during gestation and lactation, with a control group included for comparison. The offspring were subjected to behavioral assessments, including open field, marble burying, and jump tests, to evaluate locomotor activity, attention, and impulsivity. Neurochemical analyses were conducted to measure the expression of tyrosine hydroxylase (TH), aromatic L-amino acid decarboxylase (DDC), and dopamine receptor D1 (Drd1) in the PFC using Western blotting and quantitative PCR (qPCR). The results indicated that high-dose PFAS exposure (10 mg/L) resulted in hyperactivity, attention deficits, and impulsive behavior (*p* < 0.05), along with decreased TH and DDC expression and altered Drd1 levels, suggesting disrupted dopamine metabolism. These findings imply that perinatal PFAS exposure leads to behavioral abnormalities and disruptions in dopamine-related processes.

## 1. Introduction

PFASs are a class of persistent organic compounds with stable properties that emerged in the 1930s. Due to the substitution of fluorine atoms for hydrogen atoms, PFASs form covalent bonds with high bond energy, resulting in exceptional stability, water resistance, and oil repellency ^[1]^. PFASs have been detected globally owing to their persistence and have numerous industrial applications, including the production of cleaning agents, paper coatings, and food packaging materials ^[2]^. Consequently, PFASs have become a pervasive global environmental pollutant [3, 4]. After decades of production and consumption, PFASs have been widely distributed in surface water ^[5]^, soil ^[6]^, and other environments. They have also been detected in varying concentrations in human serum ^[7]^, urine ^[8]^, and breast milk ^[9]^.

PFASs in the environment can enter the human body through exposure routes such as drinking water, food, and dust. Once inside the body, these compounds tightly bind to plasma proteins and accumulate in tissues like the liver and kidneys. Due to their resistance to metabolic breakdown and excretion, PFASs pose a persistent challenge for the human body ^[10, 11]^.

Contamination of drinking water by PFASs is a global concern, with documented presence in surface and commercially available drinking water across Europe, the USA, Asia, and Australia in 2022 ^[12]^. Recent studies have identified significant levels of PFASs in drinking water sources in several developing countries, affecting both surface and groundwater systems ^[13]^. A study in the Middle East reported PFAS concentrations up to 956 ng/L in coastal waters of Saudi Arabia ^[14]^, while similar contamination has been observed in Turkey ^[15]^, and various South American and Caribbean countries ^[12]^. In China, a 2017 survey revealed that per capita exposure to PFASs in drinking water was 2-5 times higher than in developed nations ^[16]^. Contamination was also reported in Mediterranean rivers in 2016 ^[17]^, as well as in the Llobregat and Jucar rivers in Spain ^[17, 18]^. Furthermore, PFASs were detected in human serum and breast milk of 419 pregnant women in Lebanon, potentially linked to exposure through contaminated drinking water ^[19]^. Environmental toxicology research indicates that PFASs, as recognized potential human carcinogens, can cause adverse health effects such as reproductive damage ^[20]^, thyroid endocrine abnormalities ^[21]^, immunotoxicity ^[22]^, and may even exacerbate the severity of COVID-19 infections ^[23]^. These findings underscore the urgent need for attention and action.

Given the ubiquity of PFASs in the environment and their high detection rates in the human body, epidemiological studies on PFAS pollution and related diseases have been conducted. While it has been established that PFASs exhibit hepatotoxicity and can inhibit immune cell activity and cytokine release, the specific biological processes and pathogenic mechanisms remain largely unclear.

Attention deficit hyperactivity disorder (ADHD), also referred to as hyperkinetic disorder, is a persistent neurodevelopmental condition characterized by age-inappropriate levels of inattention, hyperactivity, and impulsivity ^[24]^, affecting approximately 8-12% of children worldwide ^[25]^. It is one of the most prevalent neurodevelopmental disorders in childhood, with symptoms persisting into adulthood for some individuals. As of 2023, the global prevalence of ADHD among children and adolescents is estimated to be between 5 and 7 percent ^[26]^. ADHD arises from the interplay of genetic and environmental factors. The most widely accepted hypothesis regarding its pathogenesis is the monoamine transmitter deficiency theory, which posits an imbalance of dopamine and norepinephrine in the brain ^[24]^.

A study conducted in the Dutch Maternal and Infant Cohort (N=59) revealed that prenatal exposure to PFASs was associated with externalizing problem behaviors, including hyperactivity and destructive or aggressive behaviors, in 18-month-old infants ^[27]^. A previous evaluation of ADHD-related neurobehavioral symptoms in Taiwan’s prenatal exposure cohort (N=282) at age 7 found a significant association between PFAS exposure and symptoms such as inattention and oppositional defiant disorder ^[28]^. Meanwhile, a Danish study examining children diagnosed with ADHD (N=220) reported that maternal plasma PFAS levels during pregnancy were higher in mothers of ADHD children compared to the healthy control group, suggesting a correlation between prenatal PFAS exposure and ADHD development ^[29]^. These studies collectively indicate a potential link between perinatal PFAS exposure and ADHD. However, while epidemiological evidence from three cohort studies in the Netherlands, Taiwan, and Denmark supports this association, further research is needed to establish a more robust epidemiological foundation and explore the underlying mechanisms.

In conclusion, PFASs demonstrate neurotoxic properties, and perinatal exposure to these compounds adversely affects the development of the nervous system in children, potentially contributing to the onset and progression of ADHD. Therefore, this study designed an experimental model using Sprague-Dawley (SD) rats exposed to aldehyde-fluoride compounds at varying concentrations during the perinatal period. Through a series of animal behavior experiments, the effects of these compounds on behavioral outcomes were examined, thereby elucidating the potential mechanisms underlying ADHD-like behaviors. This study, which aims to understand the impact of environmental pollutants on children’s neurodevelopment, provides new insights and highlights the importance of controlling exposure to PFASs.

## 2. Methods

### 2.1. Animal and behavioral experiments

#### 2.1.1 Animals

Male SD rats were purchased from Beijing Weidong Lihua Laboratory Animal Co., LTD. Six-week-old male SHR and age- and sex-matched Wistar Kyoto (WKY) rats, weighing 120 to 150 grams, were provided by Shanghai Slack Laboratory Animal Co., LTD. (Shanghai, China). SHR is widely recognized as the most commonly used animal model for studying ADHD at this age, as it does not develop hypertension and exhibits all behavioral characteristics of ADHD [30, 31]. WKY rats serve as the closest genetic control for SHR. All experiments were conducted in accordance with the “Guidelines for the Care and Use of Laboratory Animals” and approved by the Animal Ethics Committee of Hangzhou Medical College (Approval No. 2023-001). Rats were provided with ad libitum access to food and water and gradually acclimated to laboratory conditions (12-hour light-dark cycle, temperature 22 ± 1 ℃, relative humidity 55 ± 5%). To minimize animal suffering, the number of animals used was kept to the minimum necessary.

Perfluoroalkyl substances (PFASs) pollution was classified into mild, moderate, and severe levels based on concentrations of 1, 3, and 10 mg/L, respectively. For the SD-group (prenatal exposure), maternal rats were exposed to different concentrations of PFASs via drinking water from gestation day 1 for 21 days, after which they were provided with clean drinking water. The offspring were subsequently raised for an additional 42 days. In contrast, for the SD+ group (postnatal exposure), neonatal rats born in a clean environment were exposed to varying concentrations of PFASs through drinking water from postnatal day 1 for 42 days. Prefrontal cortex tissues were collected from both groups on postnatal day 42(**Fig. 1**)

**Fig. 1.**
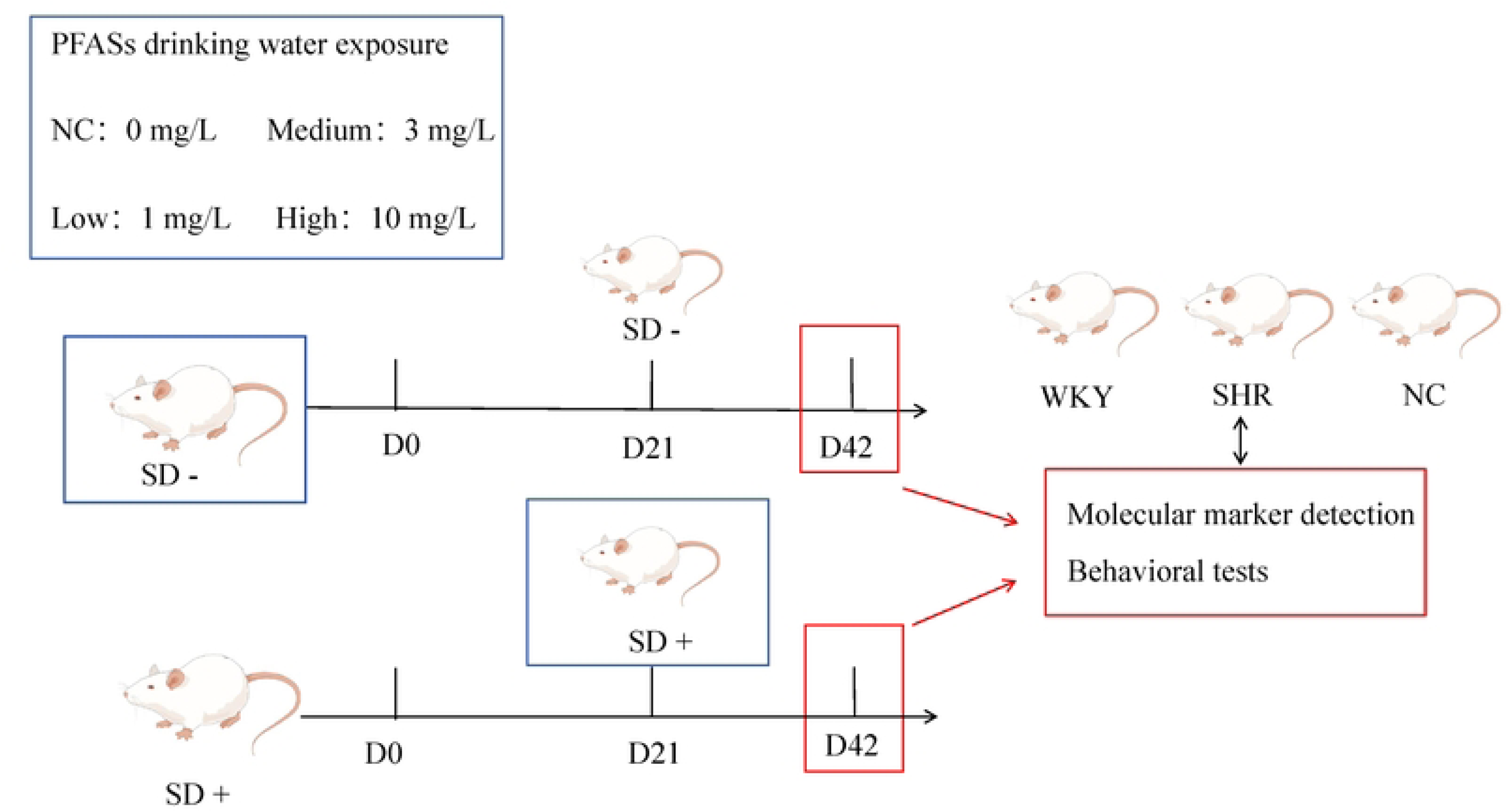
Experimental design.

### 2.2 Behavior Experiment

#### 2.2.1 Open Field Test (OFT)

The dimensions of the OFT test chamber are 100 cm × 100 cm × 40 cm. The floor is evenly divided into 16 compartments, with the central area comprising four central compartments and the peripheral area consisting of the remaining 12 compartments. At the beginning of the experiment, rats were placed in the lower left corner of the open field and allowed to explore freely for five minutes. The system automatically recorded various parameters, including the duration of movement within the central area, the frequency of entries into this region, the total distance traveled over five minutes, and the number of rearing events. The total walking distance and time spent in the central region reflect the motor activity of the rats, while the frequency of rearing indicates their exploratory behavior and interest in the environment.

#### 2.2.2 Jump Test (SDT)

Training according to Rezayof ^[32]^ was conducted in a 30 × 30 × 40 cm box, with the bottom composed of parallel stainless steel bars (0.3 cm diameter, 1 cm spacing). A wooden block measuring 4 × 4 × 4 cm was positioned in one corner of the floor. The animals were initially placed on the grid floor for 3 minutes, followed by 5 minutes of intermittent electric shocks (50 Hz, 0.3 mA, 48 V) delivered through the grid floor. After the shock period, the number of times the rats jumped off the block within 5 minutes was recorded. Twenty-four hours after training, the animals were again placed on the wooden block, and the latency from standing on the block to jumping off with all four paws on the grid floor was measured as the incubation time. The frequency of jumps from the block served as an indicator of learning performance, reflecting memory activity.

#### 2.2.3 Marble Burial (MBT)

The experiment utilized the same cage as the one used for housing the rats. The test chamber was lined with a 5 cm layer of filler material. Glass marbles were carefully placed on top of the filler and arranged in a 4 × 6 grid pattern. At the start of the experiment, the rats were gently placed in one corner of the cage, ensuring they did not come into contact with the glass marbles. The observer then exited the laboratory. The rats were allowed to move freely within the cage for 30 minutes without access to water or food. After 30 minutes, the rats were carefully removed from the cage, taking care not to disturb the glass marbles. The number of marbles buried by the rats (with at least half of the marble’s surface area covered) was recorded. The quantity of buried marbles served as an indicator of the rat’s digging behavior, with a higher number of buried marbles reflecting a greater degree of digging activity.

### 2.3 Reverse transcription polymerase chain reaction

Total RNA (1 μg) was reverse transcribed into cDNA using the Reverse Transcription Kit (EZBiosicience) and subsequently amplified by qRT-PCR using Bestar SYBR Green qPCR Master Mix (DBI Bioscience, Shanghai, China), with primers listed in Table 1 on the CFX96^TM^ Real-Time C1000 Touch System (Bio-Rad Laboratories, Hercules, CA, USA). Glyceraldehyde 3-phosphate dehydrogenase (GAPDH) served as the internal reference gene. Each PCR reaction was performed in a final volume of 20 μL, containing 2 μL of cDNA. The qRT-PCR protocol included an initial denaturation at 95 ℃ for 2 minutes, followed by 40 cycles of denaturation at 95 ℃ for 10 seconds, annealing at 58 ℃ for 30 seconds, and extension at 72 ℃ for 30 seconds. The specificity of the qRT-PCR products was confirmed by melting curve analysis. All reactions were performed in triplicate. Relative gene expression levels were normalized to GAPDH using the 2-ct method.

**Table 1.**
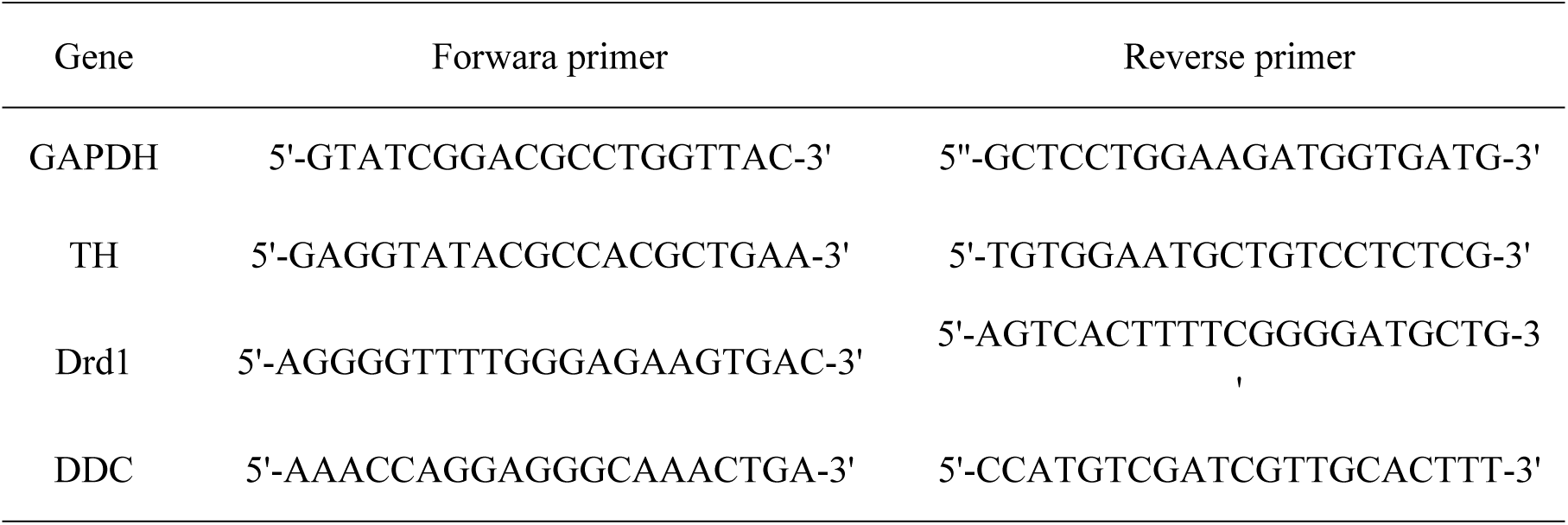
Primers for qRT-PCR analysis.

### 2.4 Western blot

Total protein was extracted from hippocampal tissues using radioimmunoprecipitation assay (RIPA) lysis buffer containing the protease inhibitor phenylmethylsulfonyl fluoride (PMSF), followed by denaturation at 95 ℃ for 5 minutes. Equal amounts of protein (30 μg) were separated by electrophoresis on 12% sodium dodecyl sulfate-polyacrylamide gels and then transferred to a polyvinylidene difluoride membrane (EMD Millipore Corporation, Billerica, MA, USA). The membrane was blocked with a quick blocking solution for 15 minutes and subsequently incubated overnight (> 12 hours) at 4 ℃ with primary antibodies against GAPDH (dilution, 1:10,000; Proteintech, Rosemont, IL, USA), TH (dilution, 1:10,000; Proteintech, Rosemont, IL, USA), and Drd1 (dilution, 1:10,000; Proteintech, Rosemont, IL, USA).

### 2.5 Statistical Methods

All experiments were performed independently a minimum of three times. One-way analysis of variance was employed for comparisons among groups. Data analysis was conducted using GraphPad statistical software. Prior to group comparisons, the normality of data distributions was assessed using the Shapiro-Wilk test. For normally distributed data, differences between groups were evaluated using the unpaired t-test, while non-normally distributed data were analyzed using the Kruskal-Wallis test. A *p*-value less than 0.05 was considered statistically significant.

## 3. Results

### 3.1 High concentration exposure group (10mg/L +) exhibits ADHD-like phenotypes, including hyperactivity and attention deficits

The open field test (OFT), marble burial test (MBT), and jump test (SDT) were conducted to explore the effects of PFAS on the behavioral and cognitive functions of SDs (Fig. 2).

**Fig. 2.**
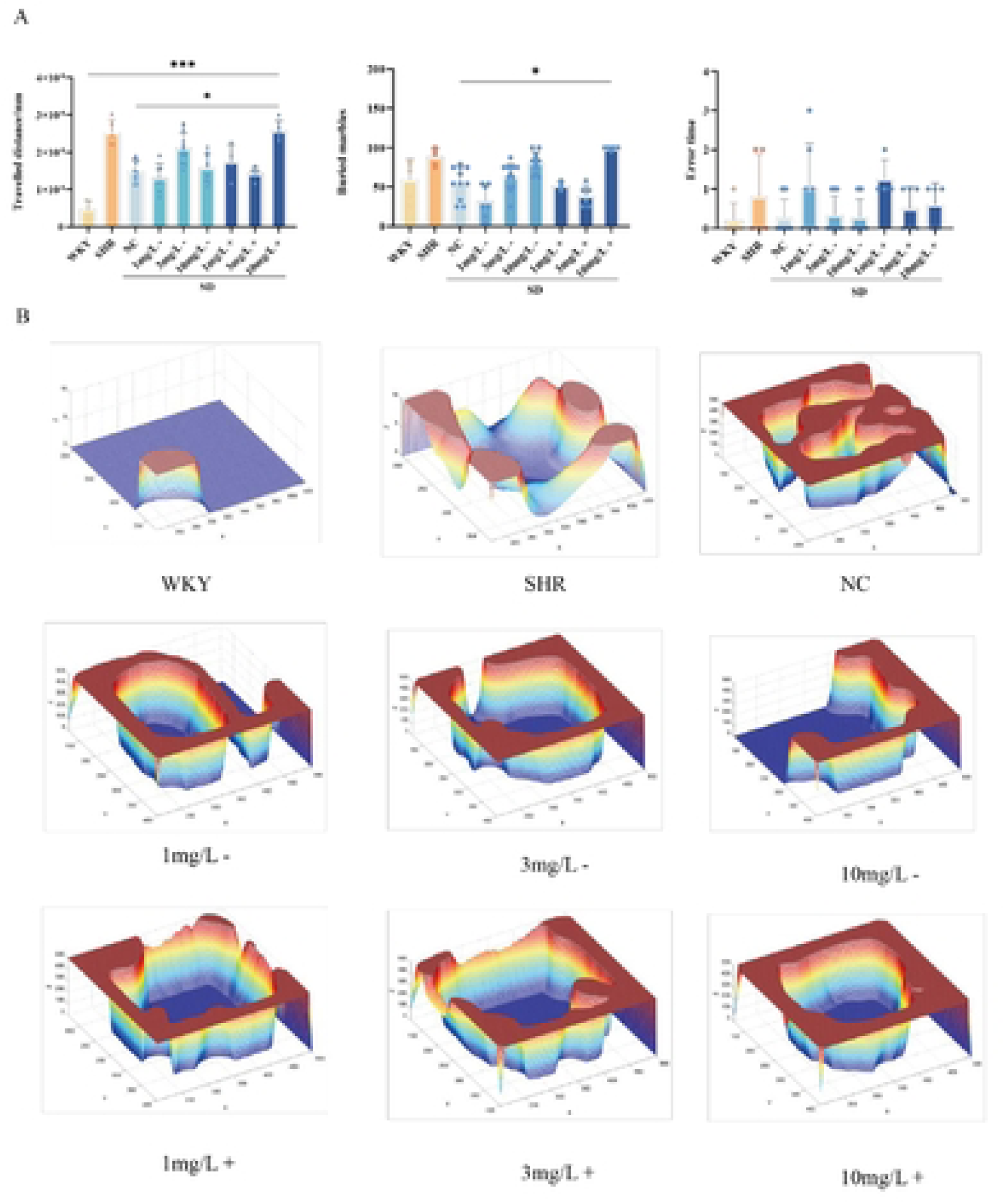
Abnormal behavior of rats exposed to PFASs. (A) The total distance traveled by each group in the open field test (OFT), the marble burial rate in the marble burial test (MBT), and the error times in the jump test (SOT). (B) 3D activity maps generated from the OFT for rats exposed to PFASs. SHR: spontaneously hypertensive rat; WKY: Wistar-Kyoto rat. Data are shown as the mean ± SEM. * p < 0.05, *** p < 0.001.

Compared with WKY and unexposed PFAS rats (NC), the high-concentration PFAS exposure group (10 mg/L +) exhibited a significantly greater total distance traveled (*p* < 0.05), and demonstrated clear signs of hyperactivity and attention deficits, such as increased spontaneous activity. In the MBT, the high-concentration PFAS exposure group (10mg/L +) showed a significantly higher burial rate (*p* < 0.05) compared to the unexposed group (NC), along with markedly increased spontaneous activity indicative of impulsivity and anxiety. In the SDT, the number of errors was significantly higher in the high-concentration PFAS exposure group compared to the unexposed group, suggesting impaired learning and memory abilities. These three sets of behavioral experiments collectively indicate that offspring exposed to high concentrations of PFAS (10mg/L) show prominent symptoms of hyperactivity, attention deficit, and other core features of ADHD, which are consistent with those observed in SHR rats with ADHD-like behavior (Fig. 2).

### 3.2 High concentration exposure group (10mg/L +) reduces dopaminergic neurotransmitter expression

#### 3.2.1 Levels of ADHD-related neurotransmitters in the PFC were measured by qPCR (Fig. 3)

In the prefrontal cortex (PFC), PCR detection revealed statistically significant differences in TH expression between the WKY group and the 10 mg/L + exposure group. Specifically, TH expression was significantly decreased in the high-dose exposure group, suggesting that PFAS exposure may inhibit dopamine synthesis. This finding is consistent with the hypothesis of insufficient dopamine in ADHD, suggesting that high doses of PFAS may affect TH expression and consequently dopamine synthesis. As a key enzyme in the dopamine synthesis pathway, DDC expression in the SHR group exposed to 3 mg/L - exposure PFAS was also examined. The results suggest that PFAS exposure may indirectly affect dopamine metabolism by altering DDC expression. Additionally, there were significant differences in Drd1 expression in the SHR group compared to the 10 mg/L - exposure group, implying that high-dose PFAS exposure may influence neuroconduction and behavioral responses in the brain by modulating dopamine receptor expression. This is closely associated with attention deficit and impulsive behavior observed in ADHD symptoms.

**Fig. 3.**
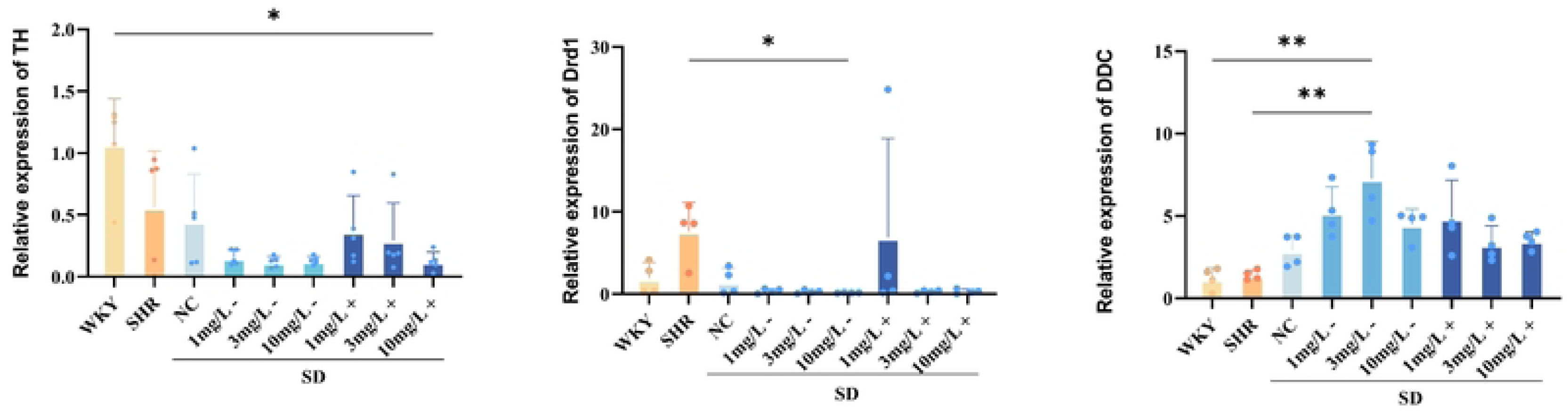
Expression of dopaminergic neurotransmitter-related genes (Drd1, TH, DOC) in the prefrontal cortex by PCR. SHR: spontaneously hypertensive rat; WKY: Wistar-Kyoto rat. Data are presented as the mean + SEM. * p < 0.05, ** p < 0.01.

#### 3.2.2 Levels of ADHD-related neurotransmitters in the PFC were quantified using Western blot (Fig. 4)

Since TH and Drd1 play important roles in neurodevelopment, the impact of perinatal exposure to aldehyde-fluoride was assessed on TH and Drd1 levels in the PFC of SD rats. Western blot analysis revealed that both TH and Drd1 expression levels were significantly reduced in the 10mg/L + exposure group compared to the SD control group.

**Fig. 4.**
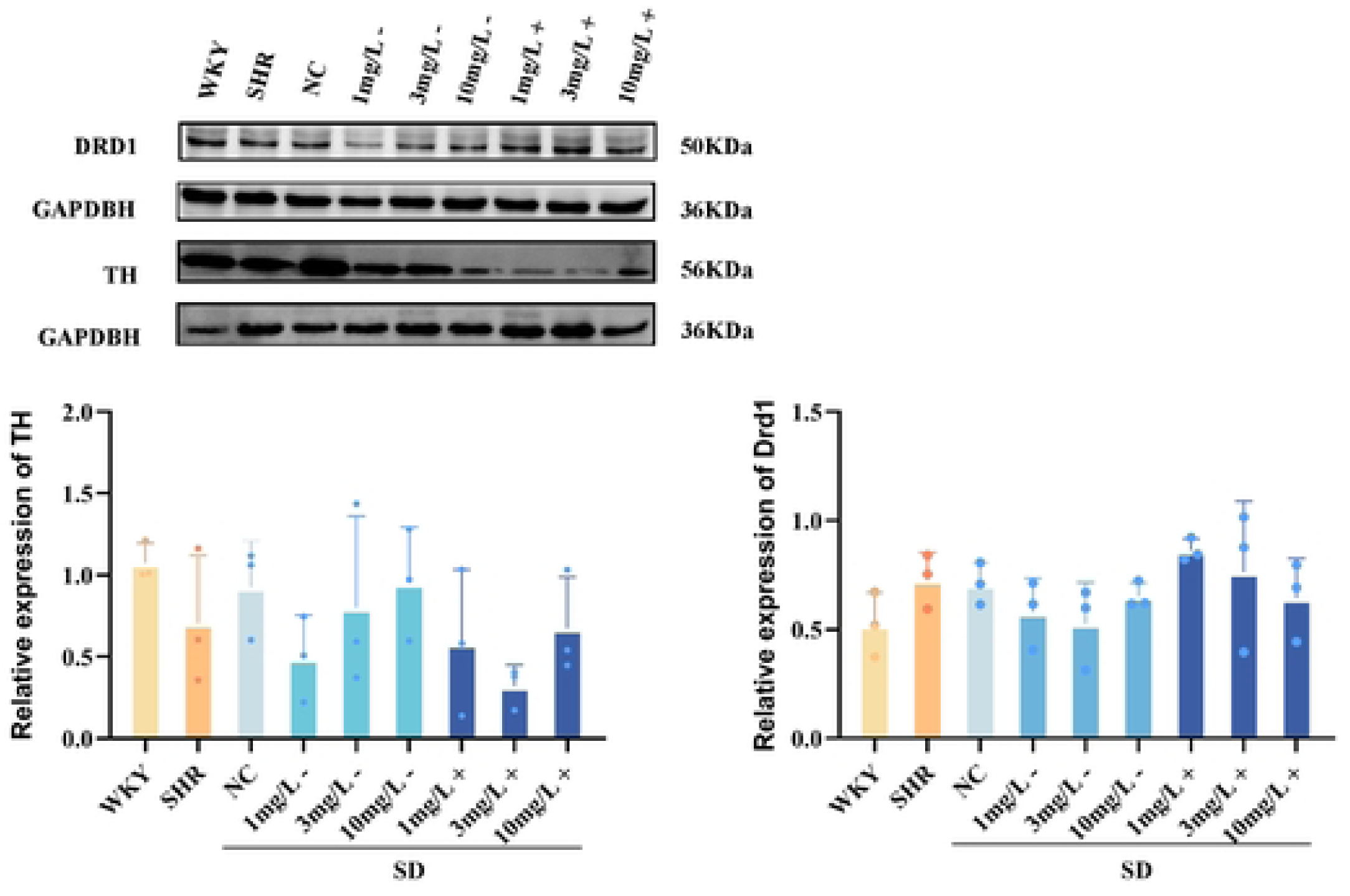
Expression of dopaminergic neurotransmitter-related proteins (TH, DRD1) in the PFC by Western blot. SHR: spontaneously hypertensive rat; WKY: Wistar-Kyoto rat.

**Fig. 4-1.**
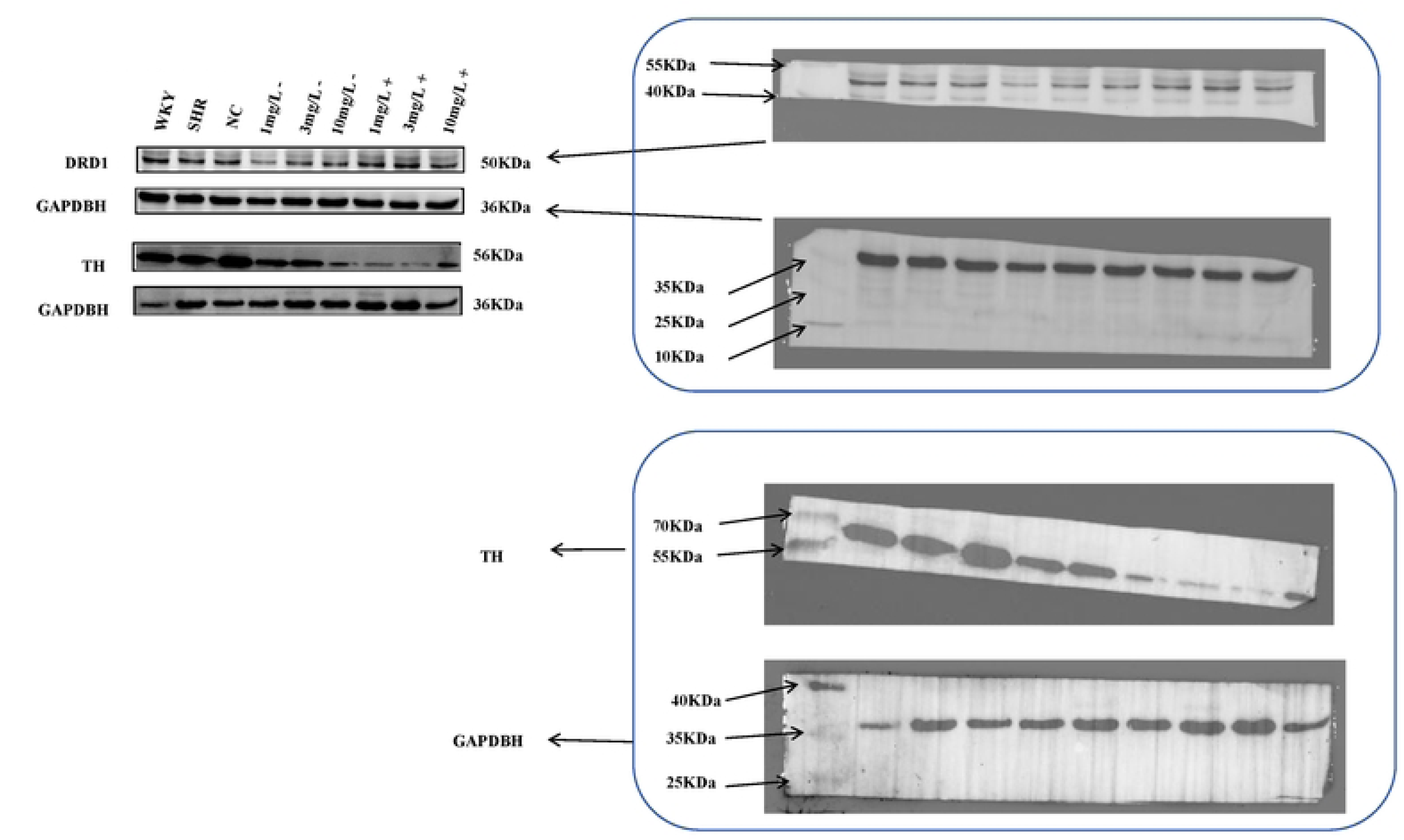
Expression of dopaminergic neurotransmitter-related proteins (TH, DRD1) in the PFC by Western blot.

**Fig. 4-2.**
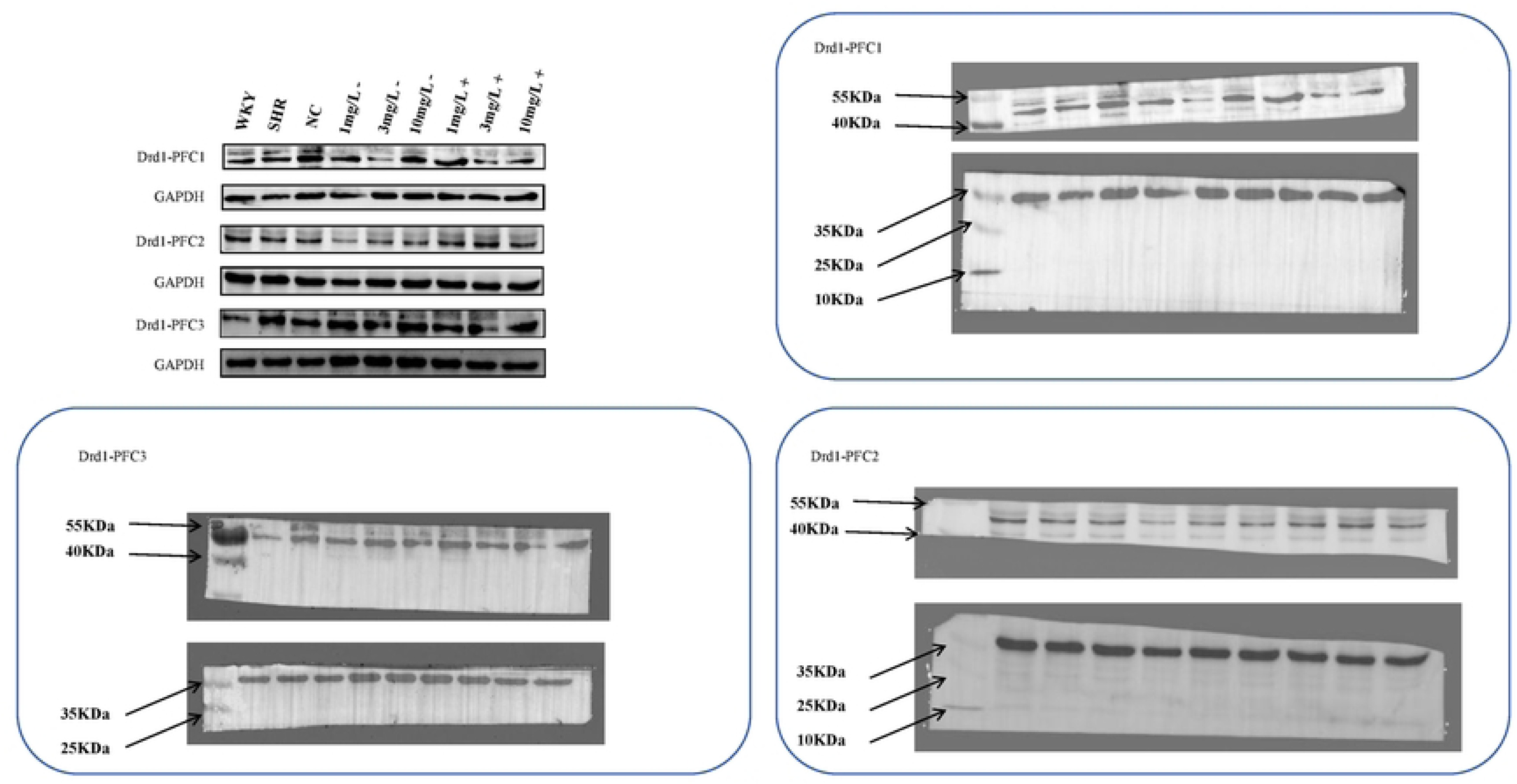
Bands involved in statistical operations of Drd I, n=3.

**Fig. 4-3.**
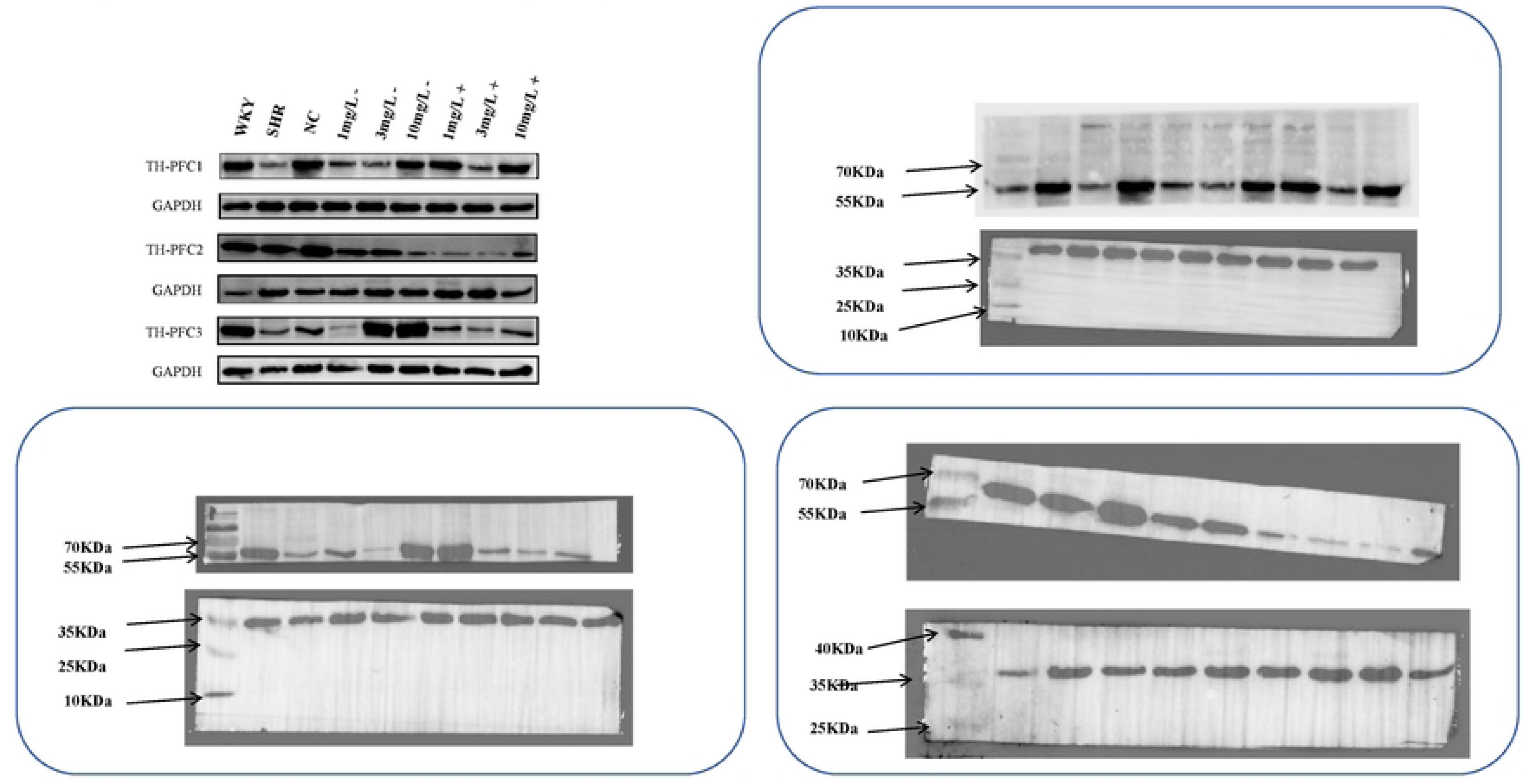
Bands involved in statistical operations of TH, n=3.

## 4. Discussion

Perfluoroalkyl substances (PFAS) are a class of ubiquitous and persistent environmental contaminants. Perfluorooctane sulfonate (PFOS), perfluorooctanoic acid (PFOA), perfluorohexane sulfonate (PFHxS), and perfluorohexanoic acid (PFHxA) are several typical representative compounds within this broad category of PFAS. They have garnered significant attention due to their chemical stability and resistance to degradation^[33]^. Given their widespread distribution in the environment and their bioaccumulative properties, it is crucial to investigate their specific impacts on biological functions, particularly neurodevelopment, and their mechanisms of action to scientifically understand and address the potential risks posed by these pollutants. This study observed the effects of perinatal PFAS exposure on SD rats and found that it could induce behaviors similar to attention deficit hyperactivity disorder (ADHD), along with significant disruptions in dopaminergic neurotransmission in the prefrontal cortex (PFC). These behavioral changes may be closely related to the downregulation of key enzymes in dopamine synthesis, such as tyrosine hydroxylase (TH), and dopamine receptors (Drd1). These findings strongly suggest that PFAS exposure could have significant effects on the human brain, especially during the perinatal period, which is the most sensitive phase of neurodevelopment[34, 35]. However, the molecular mechanisms behind these changes are not yet fully understood and require further in-depth exploration.

The perinatal period is a critical window for neurodevelopment, during which PFAS may have a lasting impact on dopaminergic circuits. Recently, a cohort study in Norway found that prenatal exposure to PFOA is associated with an increased risk of autism spectrum disorder and ADHD in children^[36]^.This finding suggests that the disruption of dopaminergic neurons by PFAS may affect the risk of ADHD and its clinical manifestations, particularly when exposed to these pollutants during the sensitive window of CNS development during gestation^[37]^.PFAS has been detected in human brain tissue samples[38–40], and similar phenomena have been observed in animal models under various environmental exposure or experimental conditions^[41]^.The potential mechanism of PFAS crossing the blood-brain barrier may be related to the discontinuity of tight junctions, which can increase the permeability of the endothelial cell membrane of human microvessels[42, 43]. At the same time, PFAS may also enter the brain through active uptake, where organic anion transporters (OAT) and organic anion transporter polypeptides (OATP) may mediate molecules with specific biochemical properties across the blood-brain barrier, and the related mechanisms have been reviewed^[44]^. Studies have shown that PFAS exposure during gestation can disrupt the migration and differentiation of midbrain dopaminergic neurons, leading to a long-term deficiency in the number of dopamine neurons^[45]^;xperiments in mice exposed to various perfluoro and polyfluoroalkyl substances on the 10th day after birth showed that exposure to these substances increased spontaneous activity and increased proteins crucial for synaptogenesis ^[46]^.This temporal specificity may explain the significant overexpression of ADHD-like behaviors in the postnatal exposure group (SD+ group) in this study. Additionally, a longitudinal study in the United States observed that increased levels of PFOA in children’s serum were associated with fewer ADHD-like behaviors in boys, while girls exhibited more ADHD-like behaviors^[47]^.Similarly, a cross-sectional study of 656 Faroese children found that elevated levels of PFOS and PFHx in the serum of 7-year-old girls were associated with behavioral problems, including ADHD-like manifestations^[48]^.These studies all indicate that the impact of PFAS on children’s behavior may have gender differences^[49]^.Future research that includes gender stratified analysis will help further clarify this dynamic process.

PFAS exposure can induce oxidative stress by generating reactive oxygen species (ROS) and depleting antioxidant defense systems such as glutathione (GSH)^[50]^.Dopamine synthesis mainly occurs in dopaminergic neurons, where the precursor tyrosine is converted to L-dopa under the action of TH, and then dopamine is generated under the action of aromatic L-amino acid decarboxylase (DDC). Studies have shown that PFOA exposure can reduce the expression of dopaminergic markers (such as TH, NFH, DAT)[37, 44], and dopamine concentration and metabolism are more sensitive to PFOS toxicity^[51]^,indicating that PFAS has a significant disruptive effect on the dopaminergic system^[52]^.Under oxidative stress, ROS can significantly inhibit TH activity by oxidizing the thiol groups of TH protein and destroying its cofactor BH4, thereby affecting dopamine synthesis^[53]^.In addition, ROS is also a key mediator of p53-induced apoptosis, which may trigger mitochondrial dysfunction and initiate apoptosis, further damaging dopaminergic neurons[54, 55].Inflammatory responses may also be involved, where PFOS exposure can activate microglia and astrocytes, release pro-inflammatory cytokines, and exacerbate neurotoxicity[56–58].The combined action of these mechanisms ultimately leads to the downregulation of key enzymes in dopamine synthesis, TH, and changes in dopamine levels ^[59]^.Further studies have shown that PFO can induce neurobehavioral changes in zebrafish larvae, with significant changes in the mRNA and protein levels of genes related to the dopamine pathway, TH, and dopamine transporter (DAT)^[60]^.Zhang et al.’s research showed that PFOS exposure inhibits TH expression in zebrafish larvae through ROS-mediated Nrf2/ARE pathway activation^[61]^.These studies are consistent with the downregulation of TH observed in the high-dose PFAS group in this study, suggesting that oxidative stress may be one of the mechanisms by which PFAS induces dopamine synthesis defects.

Epigenetic modifications (such as DNA methylation and histone acetylation) are also a potential mechanism by which PFAS disrupts the dopaminergic system. DNA methylation is a typical epigenetic modification that has been proposed as a potential mechanism for adverse effects on fetal health[62, 63].Several epidemiological studies have explored the association between PFAS and epigenetic changes in newborns, children, and adolescents^[64]^,mainly examining DNA methylation in cord blood or peripheral white blood cells of children [65, 66].The latest research indicates that prenatal exposure to PFAS may lead to changes in the DNA methylation profile of the placenta, with the placental DNA methylation profile related to PFOA being mainly enriched in angiogenesis and neuronal signaling pathways^[67]^. Among five candidate genes (IE, CHST7, FGF13, IRS4, PHOX2A, and PLXDC1), placental DNA methylation of CHST7, IRS4, and PLXDC1 is related to PFAS exposure^[67]^. PFAS can alter the DNA methylation patterns of key genes in neurodevelopment, including genes that regulate dopamine receptors and transporters. For example, PFOS exposure in newborn mice affected the transcription of dopamine receptor D5 (DRD5) in the hippocampus and cerebral cortex and tyrosine hydroxylase (TH), leading to reduced receptor expression and overactivity^[68]^.Animal studies on memory function have shown the importance of dopamine in cognition; it has been found that in rodents, DA stimulation can improve working memory, while DA receptor blockade can impair it^[69]^,which is consistent with our step-through test results. Similarly, maternal PFAS exposure in humans is associated with differential methylation of dopaminergic signaling-related genes (such as COMT, DAT1) in offspring[70, 71],which are associated with ADHD phenotypes ^[72]^.These findings suggest that PFAS may cause an imbalance in dopamine homeostasis by epigenetically silencing gene expression.

In summary, PFAS may cause significant disruptions in the dopaminergic system and induce ADHD-like behaviors through the interactive effects of multiple mechanisms such as exposure during critical windows, oxidative stress, inflammatory responses, and epigenetic modifications. Although this study focuses on the dopamine system, the pathophysiology of ADHD also involves norepinephrine and serotonin systems^[73]^.Therefore, PFAS may also indirectly disrupt dopamine homeostasis by interfering with these interconnected neurotransmitter pathways. Future research should quantify the levels of norepinephrine and serotonin in the prefrontal cortex (PFC) to assess a broader range of neurotransmitter imbalances.

In this study, we adhered to the principle of reducing animal use to comply with ethical requirements and minimize animal suffering. However, the reduced sample size may limit the generalizability and statistical power of the results, especially when exploring the association between PFAS exposure and neurodevelopment. Therefore, future research needs to verify these findings in larger sample sizes to further clarify the impact of PFAS on neurodevelopment.

This study has laid the foundation for exploring the potential mechanisms by which PFAS affects neurodevelopment, but how PFAS interacts with specific molecular targets and genetic susceptibility still needs further clarification. In-depth research on these issues will provide a solid scientific basis for developing regulatory policies and intervention strategies related to PFAS-associated neurodevelopmental risks.

## Data availability

Data will be made available on request.

## Acknowledgements

This work was supported by grants from the National Natural Science Foundation of China (grant numbers 82171168 and 81571359) and the Construction project of Shanghai Key Laboratory of Molecular Imaging (18DZ2260400).

## Animal testing declaration

All animal studies comply with the ARRIVE (Animal Research: In Vivo Laboratory Report) guidelines. All experiments were conducted in accordance with the “Guidelines for the Care and Use of Laboratory Animals” and approved by the Animal Ethics Committee of Hangzhou Medical College, No. 2023-001.

## Author contributions

**YS:** Conceptualization, Methodology, Software, Validation, Formal analysis. Investigation, Writing - Original Draft, Writing - Review & Editing and Visualization. **DC:** Conceptualization, Methodology, Software, Validation, Formal analysis, Investigation. Writing - Original Draft, Writing - Review & Editing and Visualization.**JL:** Validation and Formal analysis, Writing - Review & Editing and Visualization. **YY:** Validationand Formal analysis. **QC**: Validation and Visualization.**MP:** Validation and Visualization.**LW:** Supervision, Project administration, Funding acquisition and Writing - Review & Editing.

